# Antibiotic Resistant *Campylobacter* spp Isolated from Live Broilers at an Abattoir in Lusaka, Zambia

**DOI:** 10.1101/2023.09.12.557498

**Authors:** John Mukupa, Kunda Ndashe, Emmanuel Kabwali, Grace Mwanza, Bernadette Mumba

## Abstract

Antibiotic resistant *Campylobacter* spp causing campylobacteriosis continue to cause challenges in treatment of the infection. Poultry remains the main source of the foodborne disease. This study was undertaken to isolate and identify antibiotic resistant *Campylobacter* spp.

A total of 160 cloacal swabs were taken from broilers at a poultry abattoir in Lusaka. The samples were analyzed using standard bacteriological test. Antimicrobial susceptibility test was conducted following Clinical and Laboratory Standard Institute guideline and three antibiotics were used ciprofloxacin, tetracycline and erythromycin.

Results indicated that *Campylobacter* spp was isolated from 18.75% of the sample population, furthermore the isolates were resistant to erythromycin and tetracycline and susceptible to ciprofloxacin. The results highlight the growing concern of poultry being a source of multi-drug resistant pathogenic bacteria.

## Introduction

*Campylobacter* spp remains the commonest bacteria known to cause foodborne illness globally. Many studies have reported that poultry is the main reservoir of *Campylobacter* spp which is commensal in the gastrointestinal tract [1-4]. The abuse of antibiotics in poultry production has contributed to the emergence of multi-drug resistant *Campylobacter* spp [5-7].

Scientific reports in the past decade have highlighted the increase in the global incidence of campylobacteriosis [8-10]. Cases of the disease have been reported in North America, Europe and Australia. Information still remains limited on the number of countries in Africa reporting Campylobacter infection in both poultry and humans [6, 9, 11].

Antibiotics have been used successfully in poultry production for different purposes such as growth promotion, preventative or curative therapy. However, their use in poultry production has resulted in increased bacterial resistance to many antibiotics [12, 13]. Resistance of bacteria such as *Campylobacter* spp to some classes of antibiotics such as quinolones and macrolides is of great concern because of their significance in human medicine [14, 15]. Payot and colleagues (2002) reported multi-drug resistant Campylobacter spp broilers, turkeys and layers in Netherlands [16]. In South Africa, Bester and Essack (2008) revealed that *Campylobacter jejuni* and *Campylobacter coli* isolates were resistant to fluoroquinolones, tetracycline, and erythromycin [17].

Based on the possible risk of poultry meat being contaminated with faecal coliforms which the public are likely to be exposed to, a study was conducted to assess the presence of antibiotic resistant *Campylobacter* spp in live poultry at an abattoir in Lusaka District, Zambia.

The present cross-sectional study was carried out in September 2019. Cloacal swabs were collected from chickens received at a poultry abattoir in Lusaka district. The swabs were analyzed at the University of Zambia, School of Veterinary Medicine Bacteriology laboratory.

The poultry abattoir was conveniently selected as it was the only one available for sampling during the study period. Convenience samples were taken and the sample size was limited by the cost of the laboratory testing, a total of 100 samples were collected for the study. Every 25th processed broiler was swabbed in the cloaca using the one swab, this was done just before being stunning and slaughter. Each swab was then placed in test tube with transport media, Cary Blair (Himedia, India) and stored in a cooler box before being transported to the laboratory.

The swabs were placed in *Campylobacter* enrichment (Bolton) broth base supplemented with *Campylobacter* selective supplement IV (Himedia, India) and 5% defibrinated sheep blood. After incubation at 42 C for 24 h in a microaerophilic environment (5% O2, 10% CO2, 85% N2), 0.1 ml was streaked onto *Campylobacter* selective agar base (Himedia, India) containing an antibiotic supplement for the selective isolation of *Campylobacter* spp (Himedia, India) and 5% (v/v) defibrinated sheep blood and incubated for 48 h at 42 degrees Celsius under the same conditions. One presumptive *Campylobacter* colony from each selective agar plate was subcultured, then Gram stained and tested for production of catalase, and oxidase.

The antimicrobial susceptibility testing was done using the Kirby-Bauer disc diffusion method on Mueller Hinton Agar (Becton, Dickinson and Company, MD, USA) based on the Clinical Laboratory Standard Institute (CLSI) guidelines [18]. The antibiotic discs (Becton, Dickinson and Company, MD, USA) used included ciprofloxacin [5 μg], erythromycin [15 μg], and tetracycline [30µg].

Interpretation of susceptibility patterns on anti-microbial discs was done using guidelines laid down in the clinical laboratory standards institute (CLSI) [18] which provide break points corresponding to zone of inhibition diameter.

The study was conducted after approval from both the Lusaka Apex Medical University Research Ethics Committee (0035-19), and the Lusaka City Council.

Of the 160 cloacal swabs, *Campylobacte*r spp was isolated from 30 [18.75%] [Table]. Overall median phenotypic results of antibiotic resistance of the Campylobacter spp isolates are presented in Figure. Multi-Drug resistance, described as resistance to erythromycin and tetracycline and susceptibility to ciprofloxacin.

**Table:** the isolation rates of Campylobacter spp from live poultry.

| Microorganism | Number of Isolates [%] |  |  |
| --- | --- | --- | --- |
|  | Positive | Negative | Total |
| *Campylobacter* spp | 30 [18.75] | 130 [81.25] | 160 [100] |

**Figure:**
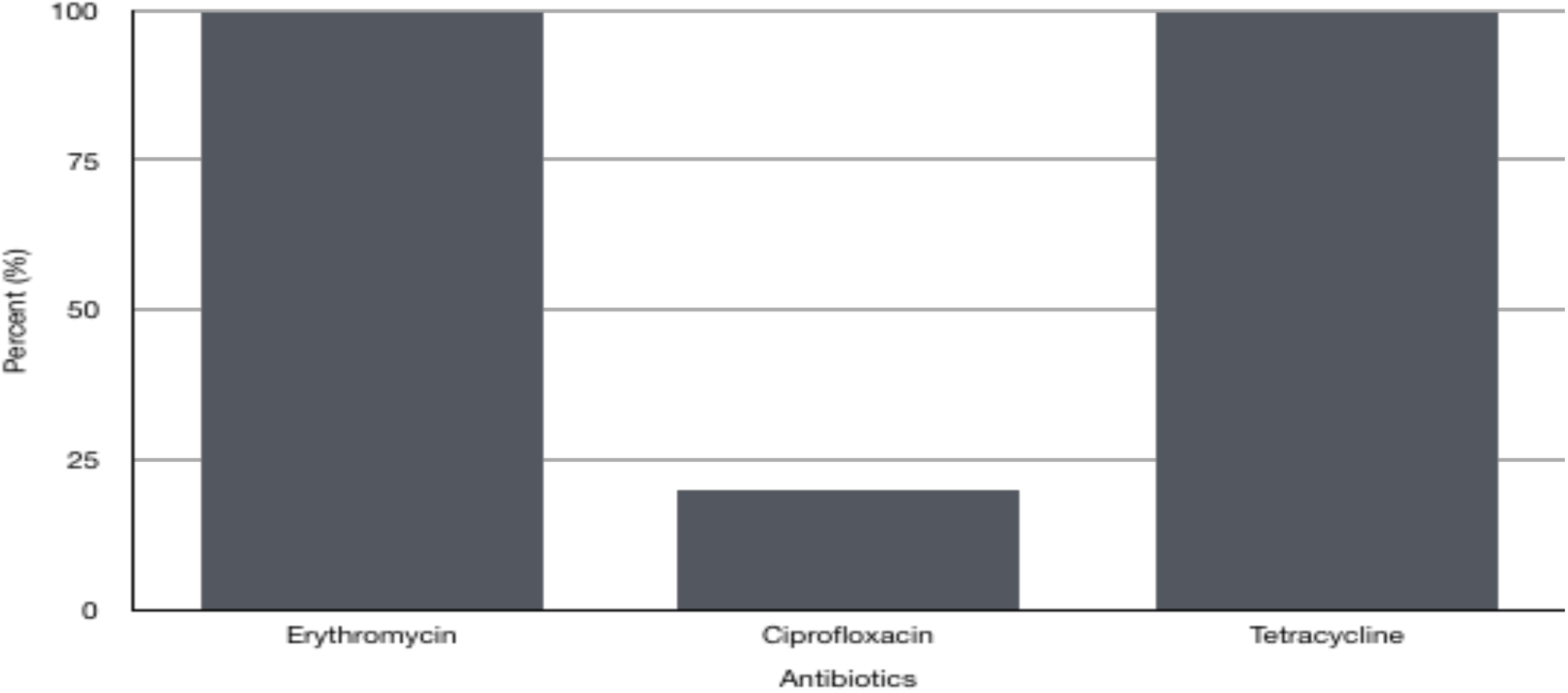
Summary data on prevalence of phenotypic resistance among in Campylobacter spp isolated from Live poultry. The antibiotics used included ciprofloxacin [5 μg], erythromycin [15 μg], and tetracycline [30µg].

In Zambia, antimicrobial medications are used poultry production for two basic reasons: (1) to treat specific infections; (2) to prevent diseases in general or to improve growth efficiency. Most concerns over the use of antimicrobials in poultry production centers on the development of antimicrobial resistance, which occurs in direct response to antimicrobial therapy in both human and veterinary medicine [13].

*Campylobacter* spp were present in 30 (18.75 %) cloacal swabs which was expected because *Campylobacter* are frequent colonizers of the intestinal tracts of birds especially poultry and falls within the reported global range of 10 to 90% [14, 19]. To the best of our knowledge this is the first study that documents the isolation rate of *Campylobacter* spp in live poultry in Lusaka, Zambia. Several studies conducted in Africa have reported particularly high prevalence of *Campylobacter* spp from from poultry. In South Africa, 47% has been reported in commercial and industrial broilers and 94% in industrial layers, 51.5% has been described in Nigeria and 63.8% in Cote d’Ivoire [17, 20-23]. *Campylobacter* spp. are common contaminants of poultry carcasses with prevalence of 20 to 100% established in fresh chicken from several countries [24-26].

In the present study resistance was observed against erythromycin and tetracycline and susceptibility to ciprofloxacin. The resistance levels in our study were comparable to work from different countries [27-30]. There are varied literature reports of resistance patterns of *Campylobacter jejuni* and *Campylobacter coli* strains; while some authors established higher resistance among *Campylobacter jejuni* [31], others found higher resistance among *Campylobacter coli* [32] and in some studies no difference in resistance were observed among the two species [33, 34].

High multi-drug resistance of *Campylobacter* spp was established in cloacal isolates which agrees with work in Malaysia by Tang et al. (2009) who recorded 100% MDR in poultry isolates [28]. Multi-drug resistances rates of 35, 75 and 97% in poultry have, respectively been reported in Malaysia, Nigeria and Thailand [27, 30]. The observed high resistance rates against most of the assayed drugs may be explained by the reality of numerous disease outbreaks which frequently threaten the Zambian poultry industry resulting in widespread use and abuse of antibiotics for prophylaxis and treatment of diseases and as growth promoters. Chicken meat is a primary source of human campylobacteriosis, therefore the presence of antimicrobial-resistant *Campylobacter* in live broilers constitutes a risk for consumers considering the high drug resistance found among isolates in this study.

*Campylobacter* spp were present in the cloacal samples of live poultry birds at the a poultry slaughter in Lusaka. Isolation and identification of multi-drug resistant Campylobacter strains poses risk to handlers and consumers. The abuse of antibiotics in poultry should be addressed due to high resistance levels documented against most of the commonly used antibiotics. Therefore, constant education, surveillance and monitoring of antibiotic usage in poultry have become necessary.

## Acknowledgement

We would like to recognize that this was a study undertaken by Mr. John Mukupa as partial fulfilment for Bachelor of Science Environmental Health of Lusaka Apex Medical University (LAMU). We wish to express our gratitude to the staff at the University of Zambia, School of Veterinary Medicine, Bacteriology Laboratory for technical help rendered during bench work sessions of the research. We could like to thank the academic staff of LAMU, faculty of Health Sciences for the support and contribution towards the research

